## Supporting information for "Separating phases of allopolyploid evolution with resynthesized and natural *Capsella bursa-pastoris*"

**Figure 4—figure supplement 1** Venn diagram of genes showed expression level dominance (ELD) of the three allotetraploid groups in leaves, separated by directions of ELD

**Figure 5—figure supplement 1** Distribution of gene homoeolog expression bias (HEB) of each allotetraploid individual in flowers and leaves

**Figure 5—figure supplement 2** The distribution of gene homoeolog expression bias of resynthesized and natural *Capsella* allotetraploids by major chromosomes in leaves

**Figure 5—figure supplement 3** Homoeolog expression bias along chromosome positions in the inflorescence sample of the Sd group

**Figure 5—figure supplement 4** Homoeolog expression bias along chromosome positions in the leaf sample of the Sd group

**Figure 5—figure supplement 5** Homoeolog expression bias along chromosome positions in the inflorescence sample of the Sh group

**Figure 5—figure supplement 6** Homoeolog expression bias along chromosome positions in the leaf sample of the Sh group

**Figure 6—figure supplement 1** Homoeolog expression bias along chromosome positions in the inflorescence sample of the Cbp group

**Figure 6—figure supplement 2** Homoeolog expression bias along chromosome positions in the leaf samples of the Cbp group

**Figure 5—figure supplement 7** Estimated number of breakpoints per chromosome quartet in resynthesized allotetraploids

**Figure 8—figure supplement 1** Relationships between homoeolog expression change and non-additive gene expression in leaves

**Figure 1–Source Data 1** *Capsella* plants used in the present study

**Figure 2—Source Data 1** Effects of plant group and positions (tray ID) on phenotypes

**Figure 4—Source Data 1** Additive and non-additive gene expression in allotetraploid groups

**Figure 8—Source Data 1** Expression level fold-change (log2FC) of homoeologs relative to the corresponding gene in diploid groups among genes with expression level dominance (ELD) in flowers or leaves

**Supplementary file 1** Tray ID of each *Capsella* plant and the position of eight plants within each tray (as a separate file)

**Supplementary file 2** RNA-sequencing information (as a separate file)

**Supplementary file 3** Differentially expressed genes (DEGs) in pair-wise contrasts among the five *Capsella* plant groups in flowers and leaves

**Supplementary file 4** Summary of differential expression analyses of allotetraploid groups

**Supplementary file 5** Gene ontology (GO) terms that were overrepresented in differentially expressed genes (FC > 1.5 and FDR < 0.05) between resynthesized and natural allopolyploids (as a separate file)

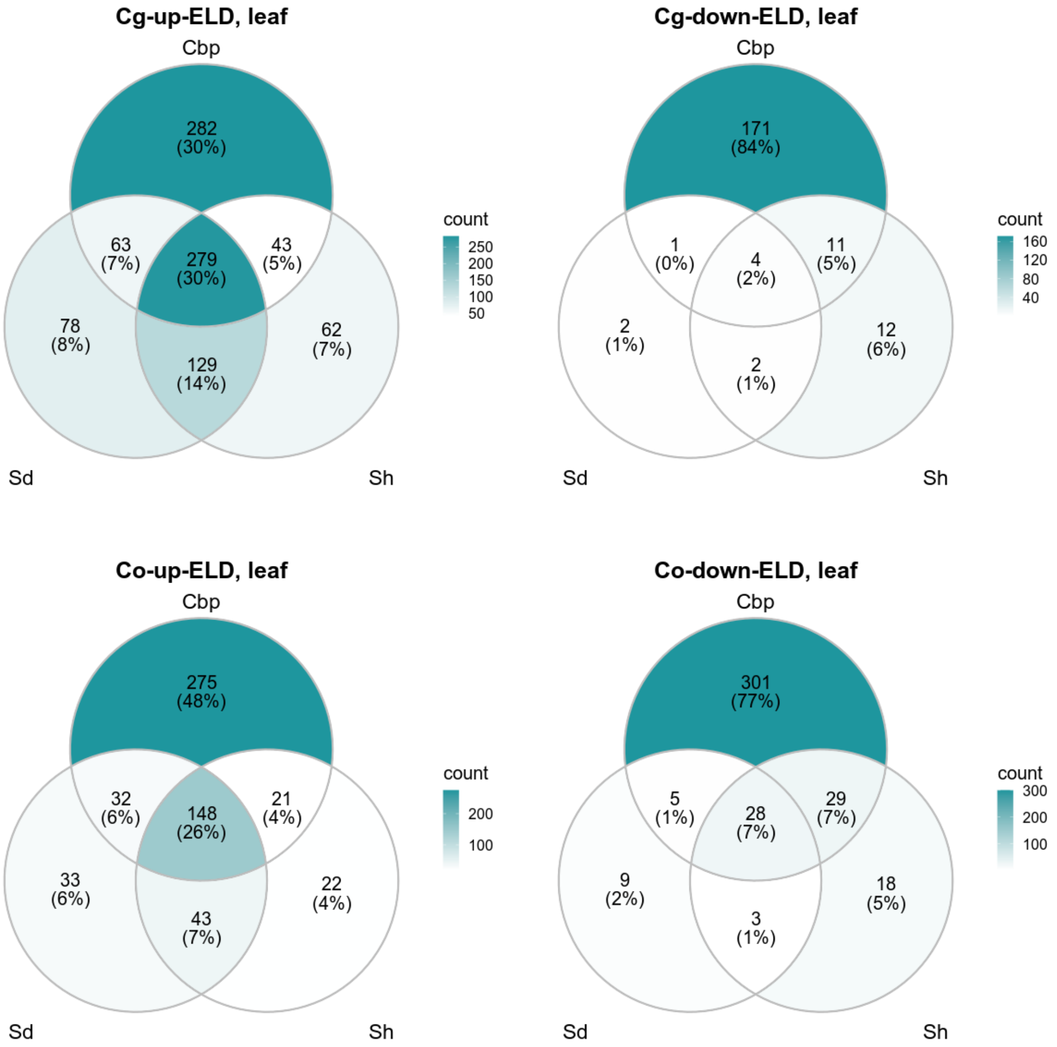


**Figure 4—figure supplement 1** Venn diagram of genes showed expression level dominance (ELD) of the three allotetraploid groups in leaves, separated by directions of ELD.

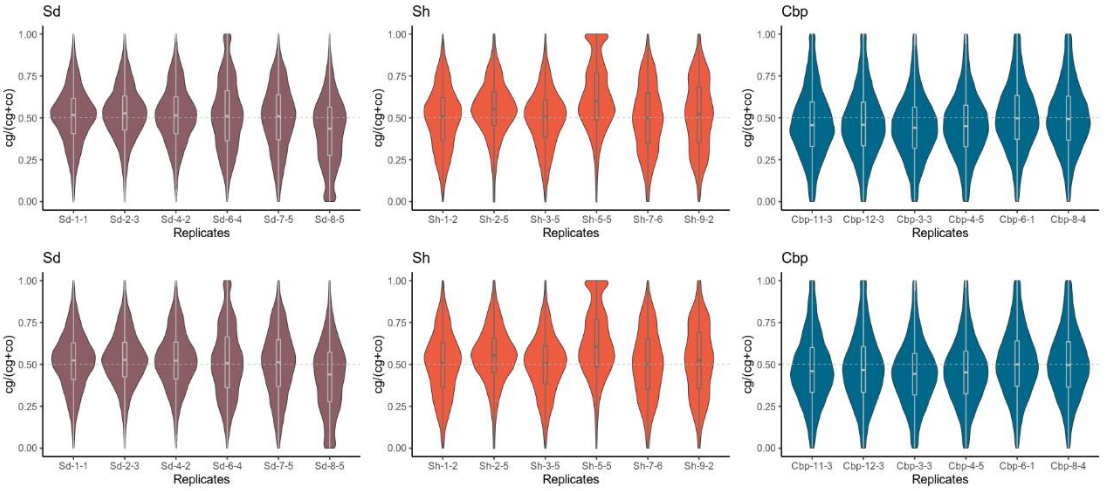


**Figure 5—figure supplement 1** Distribution of gene homoeolog expression bias (HEB) of each allotetraploid individual in flowers (upper panel) and leaves (lower panel). For each gene, HEB was estimated as the expression of *C. grandiflora*-derived homoeolog divided by the total expression of both *C. grandiflora*- and *C. orientalis*-derived homoeolog (cg/(cg+co)). HEBs were calculated for 18,255 genes in flowers, and 15,581 genes in leaves.

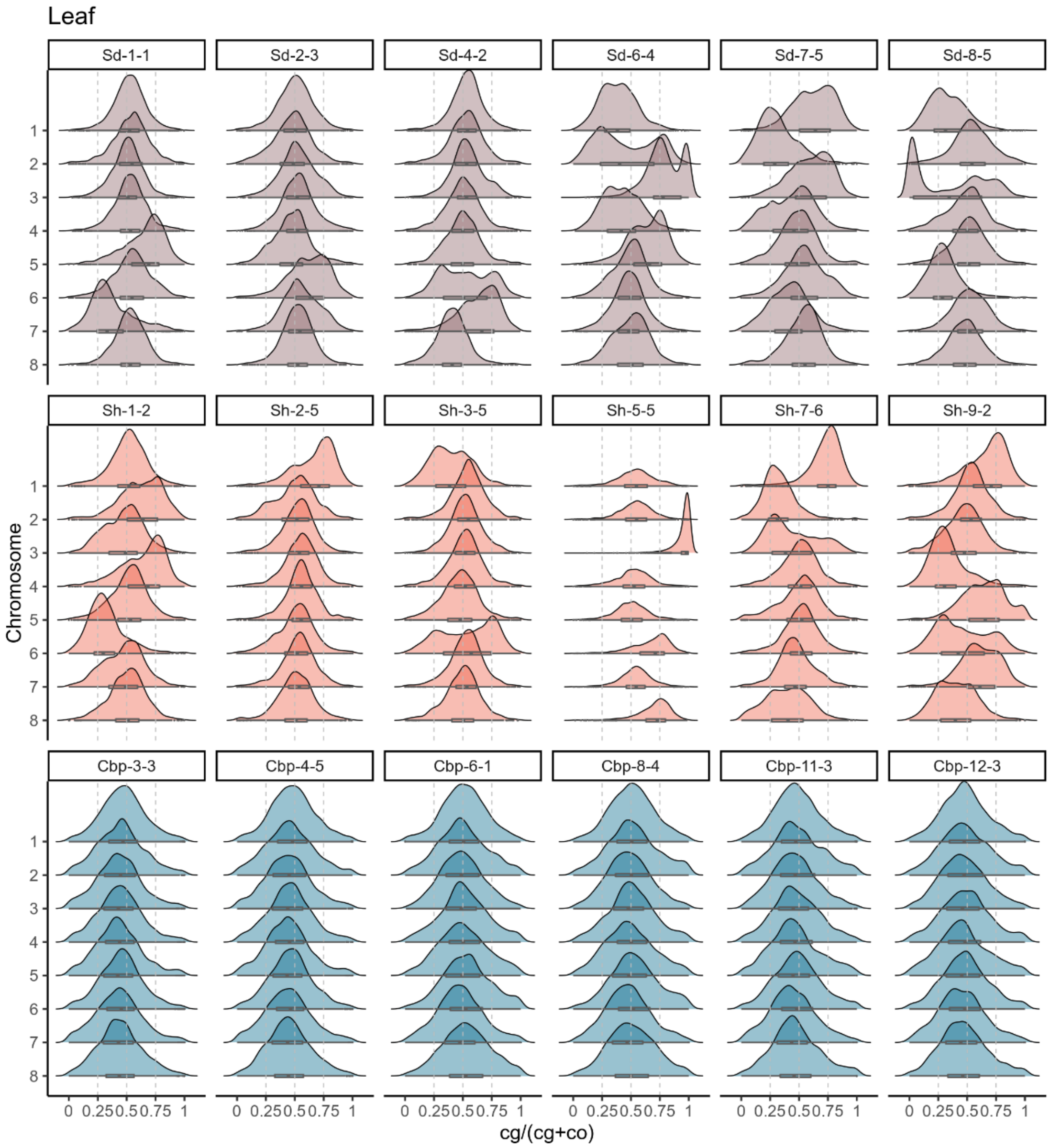


**Figure 5—figure supplement 2** The distribution of gene homoeolog expression bias of resynthesized and natural *Capsella* allotetraploids by major chromosomes in leaves.

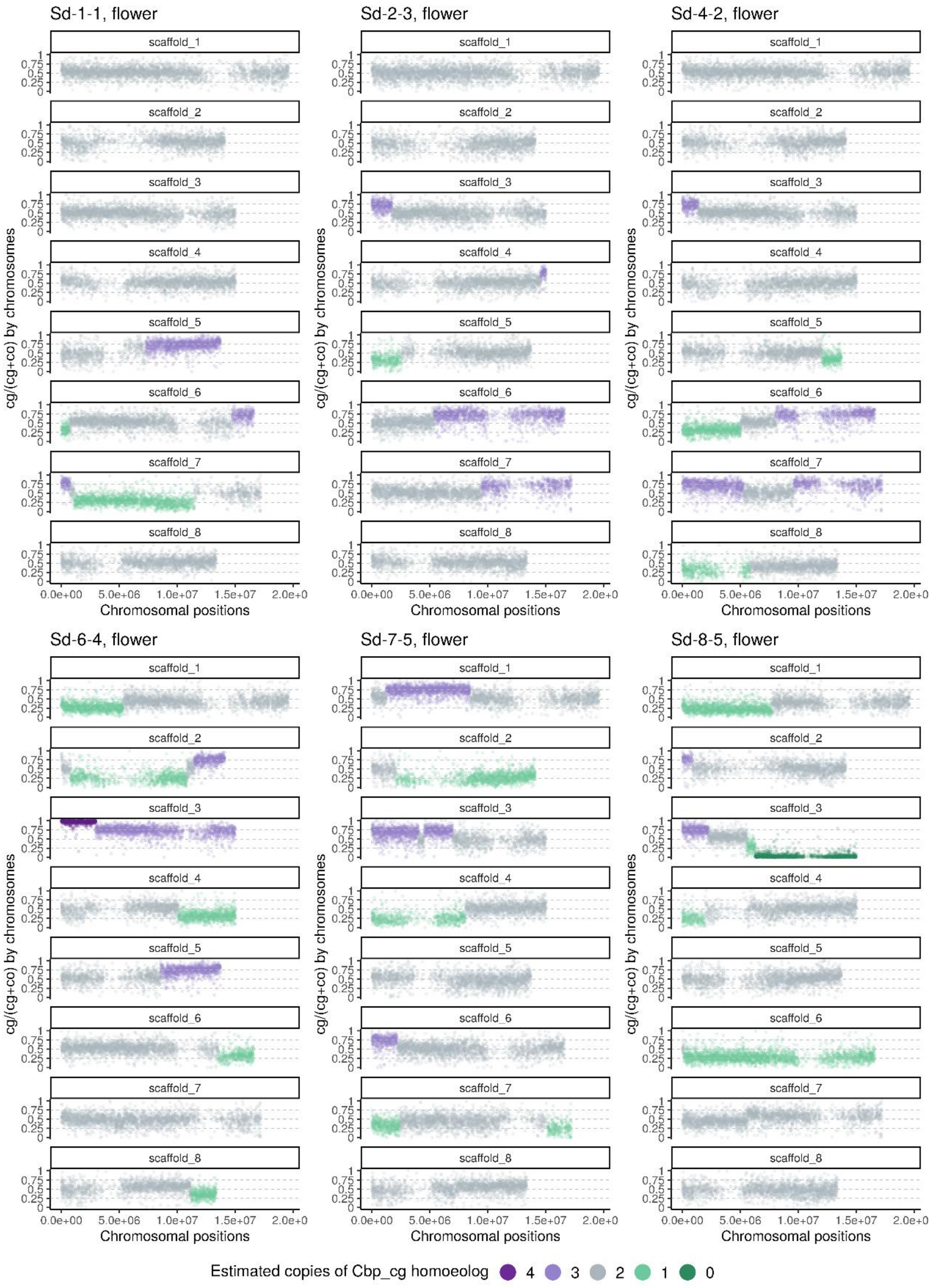


**Figure 5—figure supplement 3** Homoeolog expression bias along chromosome positions in the inflorescence sample of the Sd group ("Whole-genome-duplication-first" resynthesized *Capsella* allotetraploids). The number of cg-homoeologs at each gene estimated by the five-state Hidden Markov Model (HMM) was indicated by five colors. Dark green, light green, grey, light purple and dark purple represent (0, 1, 2, 3, 4) cg-homoeologs and (4, 3, 2, 1, 0) co-homoeologs, respectively.

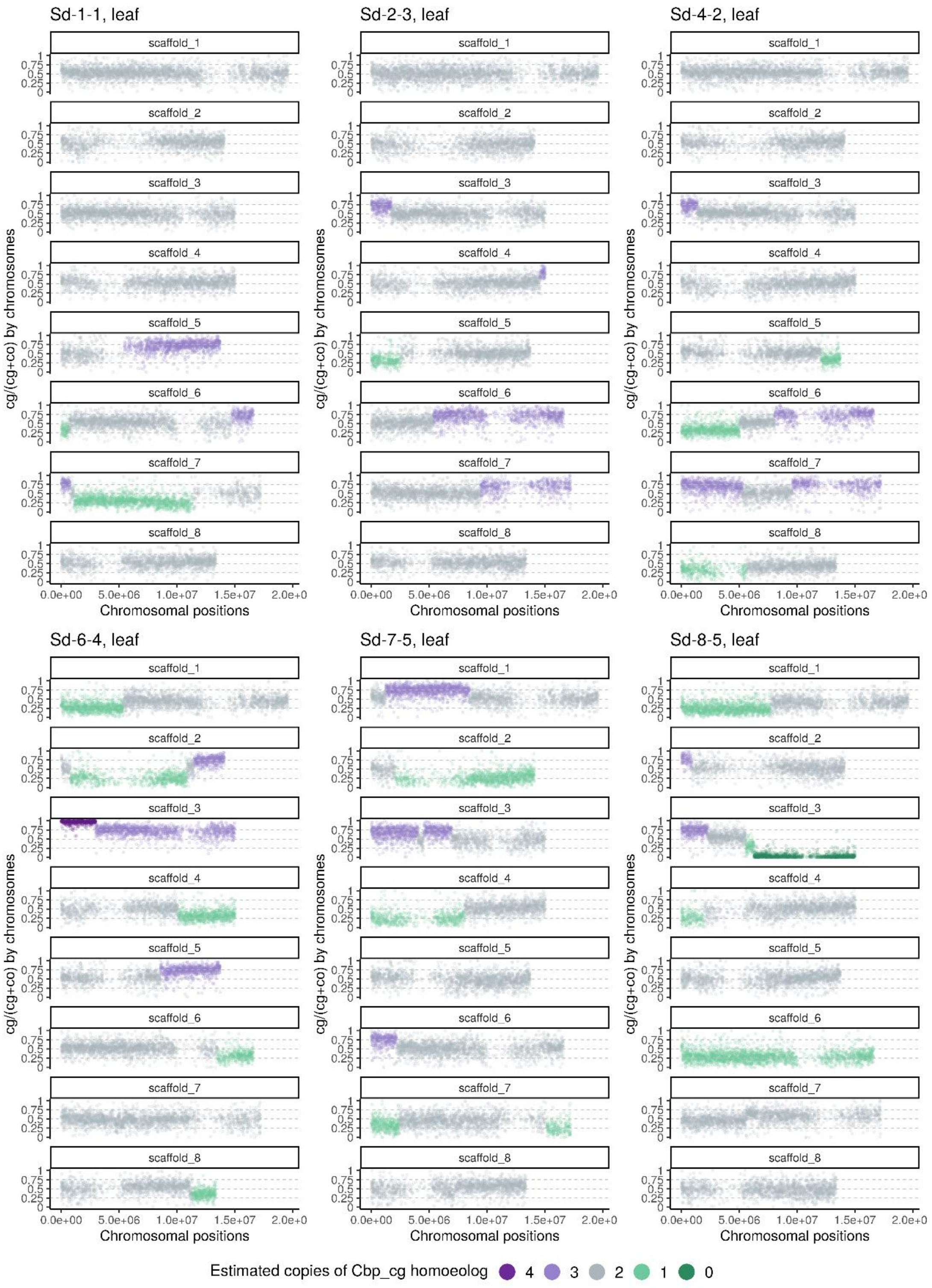


**Figure 5—figure supplement 4** Homoeolog expression bias along chromosome positions in the leaf sample of the Sd group ("Whole-genome-duplication-first" resynthesized *Capsella* allotetraploids). The number of cg-homoeologs at each gene estimated by the five-state Hidden Markov Model (HMM) was indicated by five colors. Dark green, light green, grey, light purple and dark purple represent (0, 1, 2, 3, 4) cg-homoeologs and (4, 3, 2, 1, 0) co-homoeologs, respectively.

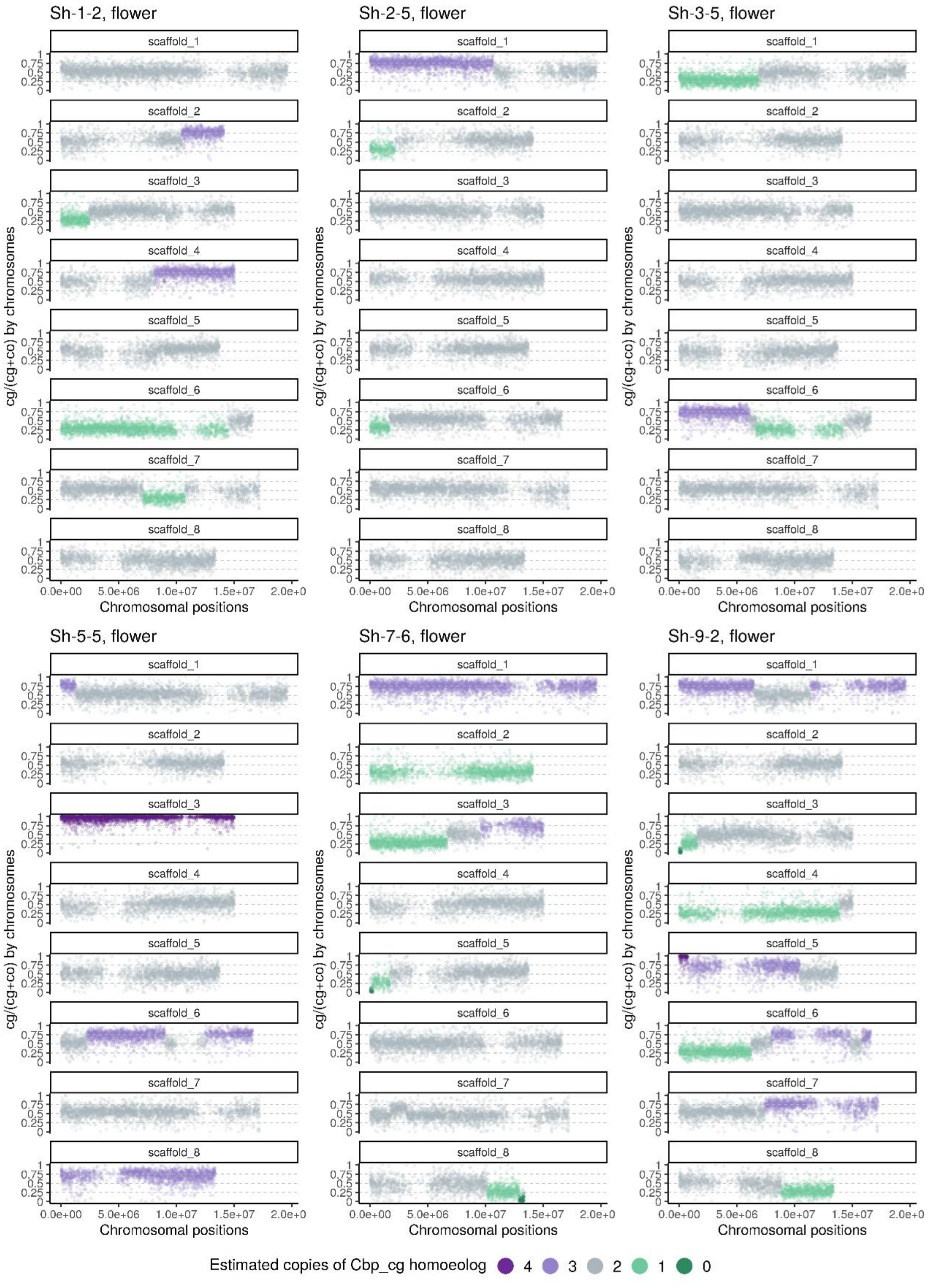


**Figure 5—figure supplement 5** Homoeolog expression bias along chromosome positions in the inflorescence sample of the Sh group (“hybridization-first” resynthesized *Capsella* allotetraploids). The number of cg-homoeologs at each gene estimated by the five-state Hidden Markov Model (HMM) was indicated by five colors. Dark green, light green, grey, light purple and dark purple represent (0, 1, 2, 3, 4) cg-homoeologs and (4, 3, 2, 1, 0) co-homoeologs, respectively.

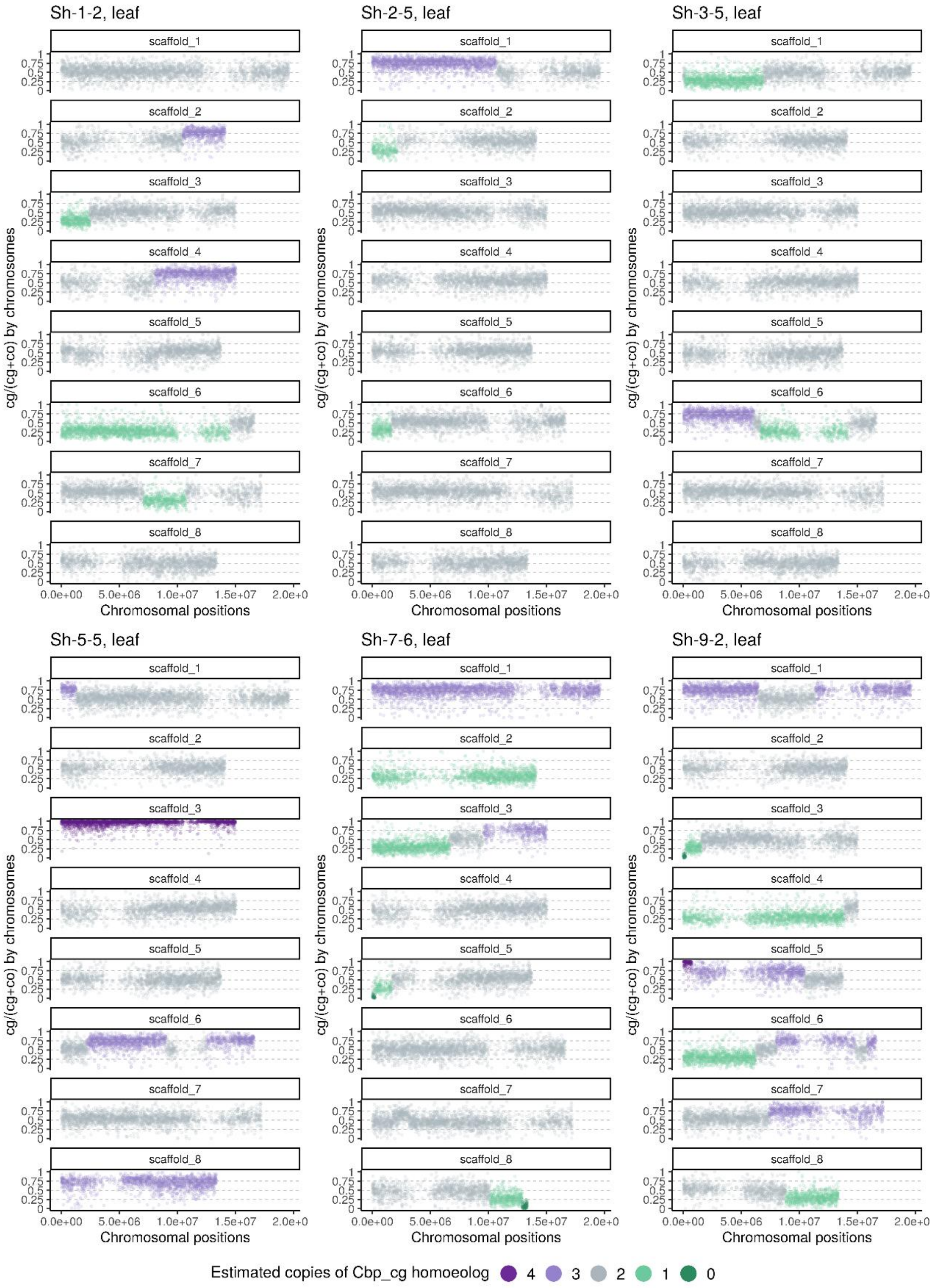


**Figure 5—figure supplement 6** Homoeolog expression bias along chromosome positions in the leaf sample of the Sh group ("hybridization-first" resynthesized *Capsella* allotetraploids). The number of cg-homoeologs at each gene estimated by the five-state Hidden Markov Model (HMM) was indicated by five colors. Dark green, light green, grey, light purple and dark purple represent (0, 1, 2, 3, 4) cg-homoeologs and (4, 3, 2, 1, 0) co-homoeologs, respectively.

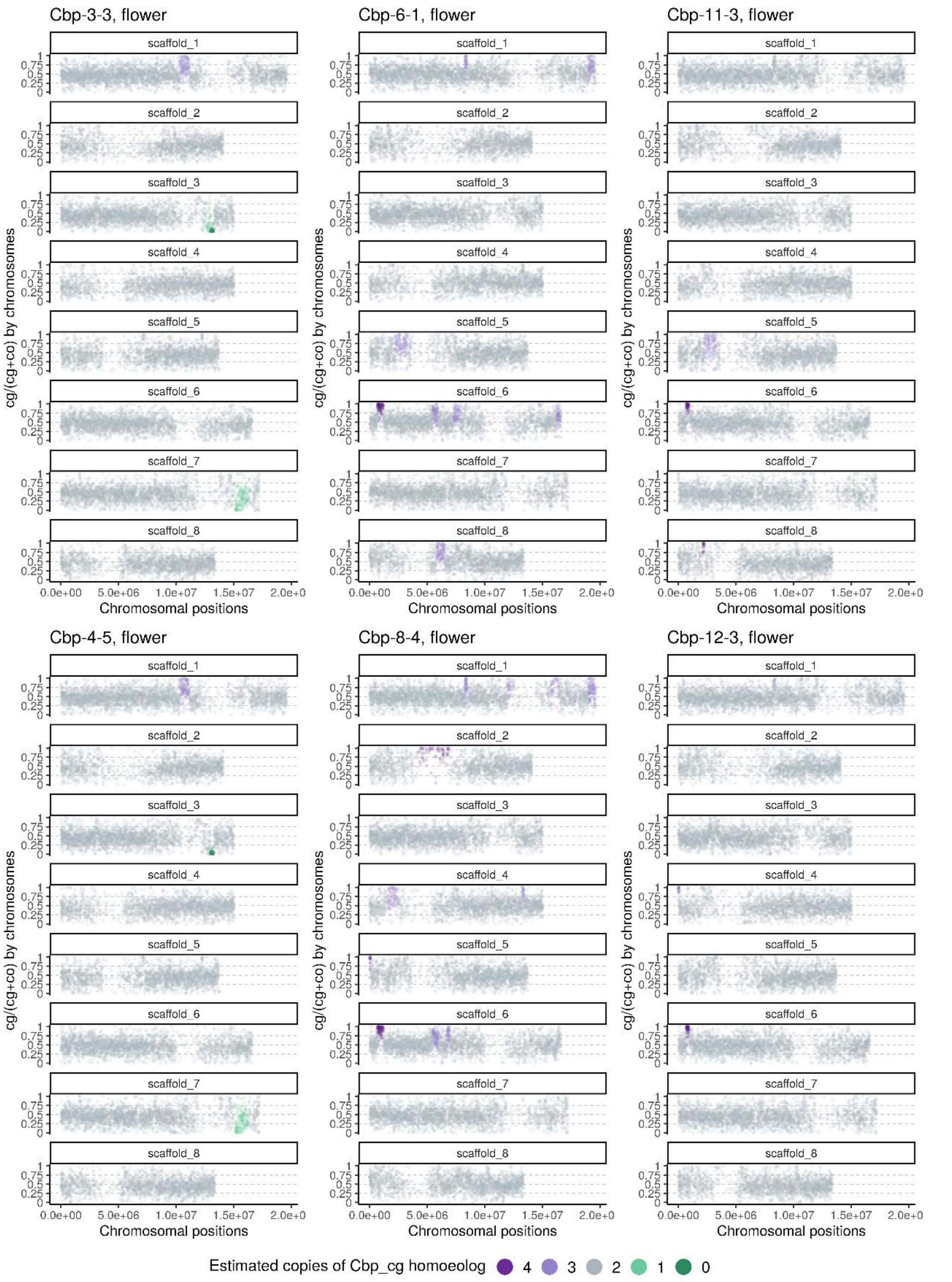


**Figure 6—figure supplement 1** Homoeolog expression bias along chromosome positions in the inflorescence sample of the Cbp group (natural *Capsella bursa-pastoris*). The number of cg-homoeologs at each gene estimated by the five-state Hidden Markov Model (HMM) was indicated by five colors. Dark green, light green, grey, light purple and dark purple represent (0, 1, 2, 3, 4) cg-homoeologs and (4, 3, 2, 1, 0) co-homoeologs, respectively.

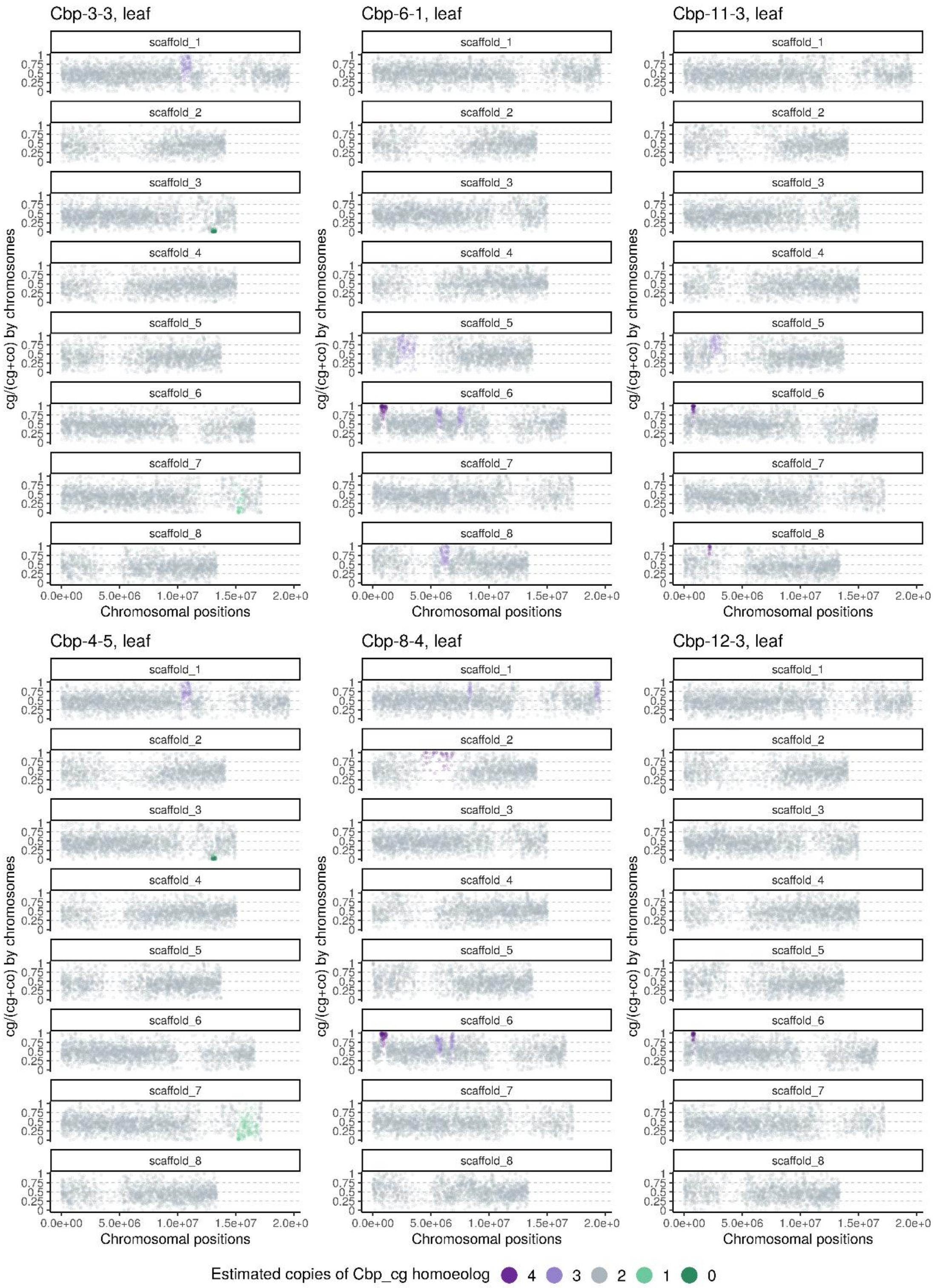


**Figure 6—figure supplement 2** Homoeolog expression bias along chromosome positions in the leaf samples of the Cbp group (natural *Capsella bursa-pastoris*). The number of cg-homoeologs at each gene estimated by the five-state Hidden Markov Model (HMM) was indicated by five colors. Dark green, light green, grey, light purple and dark purple represent (0, 1, 2, 3, 4) cg-homoeologs and (4, 3, 2, 1, 0) co-homoeologs, respectively.

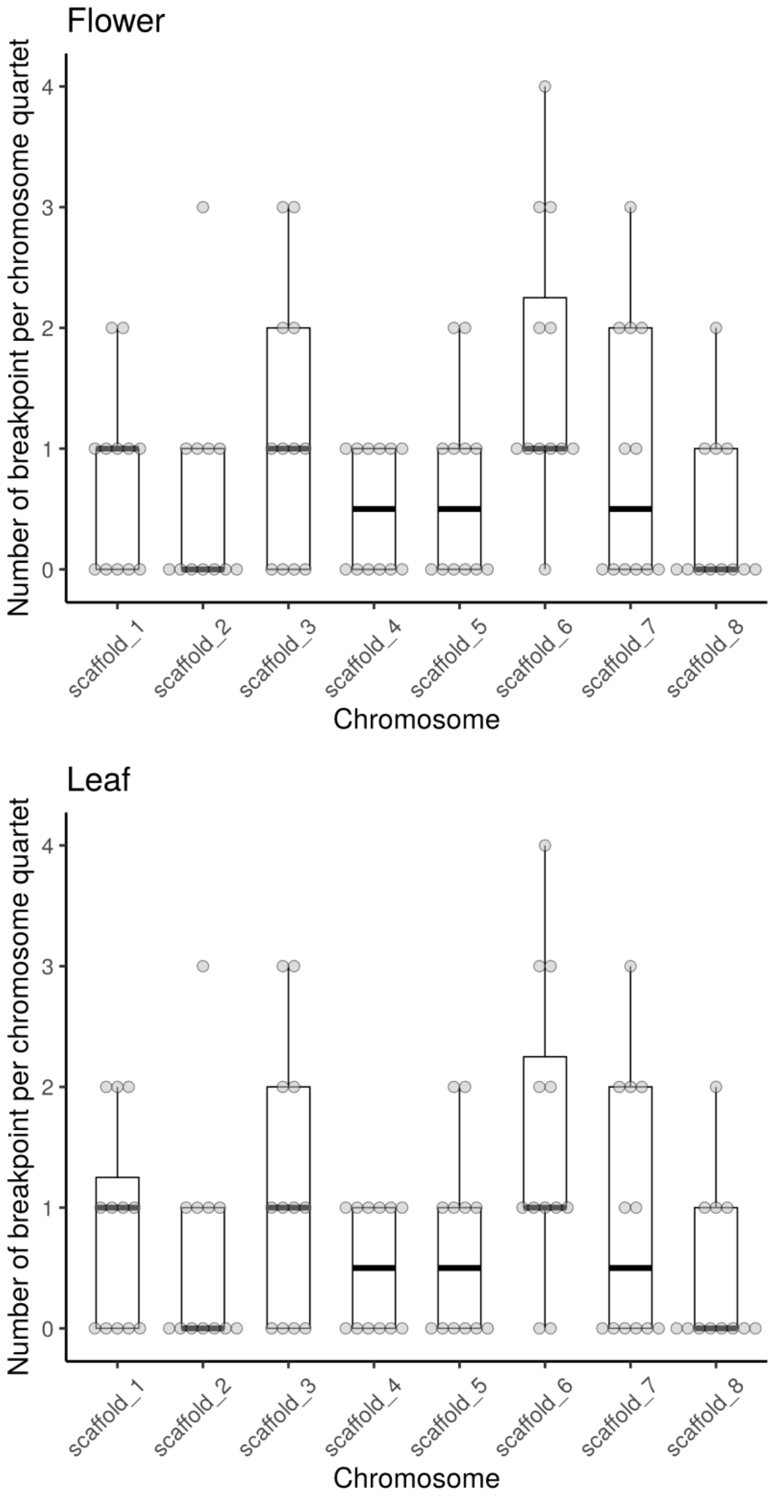


**Figure 5—figure supplement 7** Number of breakpoints between chromosome segments with distinct average homoeologous expression bias (HEB) per chromosome in flowers and leaves of the twelve resynthesized allotetraploid individuals (groups Sd and Sh), estimated with gene HEB and a five-state Hidden Markov Model. Each dot represents the estimation of one chromosome in one individual. Quartiles and median were shown by boxes and thicker bars.

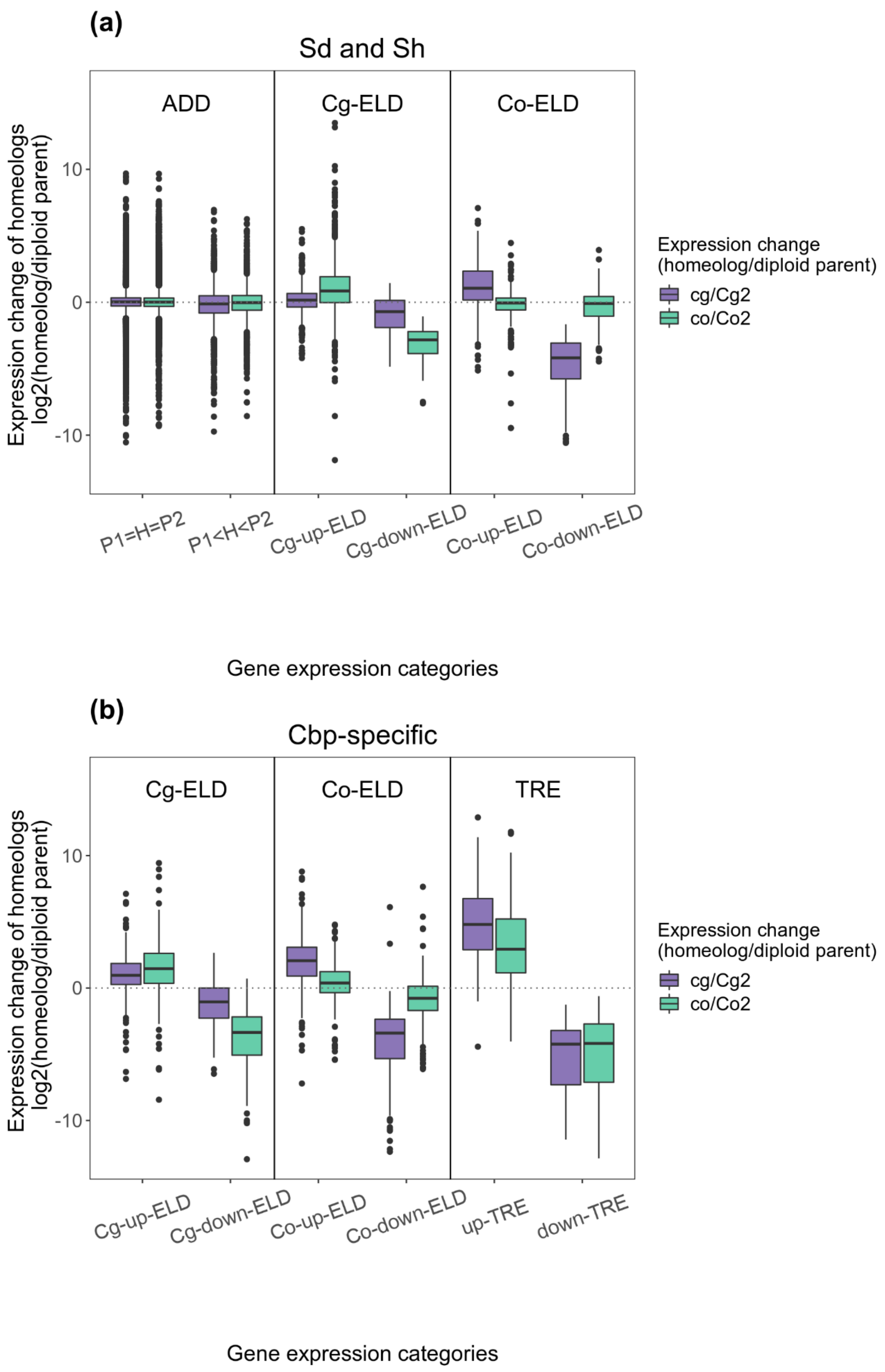


**Figure 8—figure supplement 1** Relationships between homoeolog expression change and non-additive gene expression in leaves. Gene (homoeolog) expression levels were normalized with the trimmed mean of M-values (TMM) method. (a) Homoeolog expression change among genes with ELD in resynthesized allotetraploids. (b) Homoeolog expression change among genes with Cbp-specific non-additive expression.

**Figure 1–Source Data 1** *Capsella* plants used in the present study

| **Species** | **Line ID** | **Line notation** | **Latitude (N)** | **Longitude (E)** | **Genetic cluster*** | **Reference** |
|---|---|---|---|---|---|---|
| | BEL5 | Cbp-6 | 50.55 | 128.28 | EUR | this study |
| | JO56 | Cbp-11 | 31.97 | 35.98 | ME | this study |
| *Capsella bursa-pastoris* | SE14 | Cbp-8 | 62.64 | 17.94 | EUR | this study |
| | TBS195 | Cbp-3 | 33.57 | 107.45 | ASI | this study |
| | TR83 | Cbp-12 | 41.02 | 28.97 | ME | this study |
| | TY118 | Cbp-4 | 37.55 | 112.32 | ASI | this study |
| *Capsella grandiflora* | 81 | - | 37.30 | 22.06 | - | (Duan *et al.*, 2023) |
| *Capsella orientalis* | URAL-RUS5 | - | 55.11 | 61.39 | - | (Duan *et al.*, 2023) |

* The three major genetic clusters (populations) of natural *C. bursa-pastoris* were Asian (ASI), Middle-East (ME), and Europe (EUR), defined by Kryvokhyzha *et al.* (2016, 2019).

**Figure 2–Source Data 1** Effects of plant group and positions (tray ID) on phenotypes

| | | **General linear models, ANOVA (type III tests)** | | | | | |
|---|---|---|---|---|---|---|---|
| | | **~ Group** | | | **~ Group + Tray_ID** | | |
| | | **df** | **F** | **p-value** | **df** | **F** | **p-value** |
| Petal length | Group | 4 | 296.0 | **<2.2e-16** | 4 | 310.81 | **<2.2e-16** |
| | Tray | - | - | - | 35 | 1.78 | **0.0111** |
| | Residuals | 160 | - | - | 125 | - | - |
| Petal width | Group | 4 | 384.25 | **<2.2e-16** | 4 | 388.11 | **<2.2e-16** |
| | Tray | - | - | - | 35 | 1.12 | 0.321 |
| | Residuals | 160 | - | - | 125 | - | - |
| Sepal length | Group | 4 | 194.08 | **<2.2e-16** | 4 | 218.73 | **<2.2e-16** |
| | Tray | - | - | - | 35 | 1.65 | **0.0242** |
| | Residuals | 160 | - | - | 125 | - | - |
| Sepal width | Group | 4 | 187.15 | **<2.2e-16** | 4 | 192.74 | **<2.2e-16** |
| | Tray | - | - | - | 35 | 1.18 | 0.247 |
| | Residuals | 160 | - | - | 125 | - | - |
| Pistil length | Group | 4 | 95.35 | **<2.2e-16** | 4 | 96.24 | **<2.2e-16** |
| | Tray | - | - | - | 35 | 1.18 | 0.253 |
| | Residuals | 160 | - | - | 125 | - | - |
| Pistil width | Group | 4 | 78.47 | **<2.2e-16** | 4 | 74.67 | **<2.2e-16** |
| | Tray | - | - | - | 35 | 0.879 | 0.663 |
| | Residuals | 160 | - | - | 125 | - | - |
| Stamen length | Group | 4 | 173.99 | **<2.2e-16** | 4 | 190.61 | **<2.2e-16** |
| | Tray | - | - | - | 35 | 1.46 | 0.069 |
| | Residuals | 160 | - | - | 125 | - | - |
| Stem length | Group | 4 | 84.52 | **<2.2e-16** | 4 | 90.71 | **<2.2e-16** |
| | Tray | - | - | - | 35 | 1.33 | 0.128 |
| | Residuals | 166 | - | - | 131 | - | - |
| Flowering time | Group | 4 | 49.20 | **<2.2e-16** | 4 | 25.64 | **1.13e-15** |
| | Tray | - | - | - | 35 | 1.37 | 0.104 |
| | Residuals | 165 | - | - | | - | - |
| Pollen grains per flower | Group | 4 | 164.65 | **<2.2e-16** | 4 | 145.12 | **<2.2e-16** |
| | Tray | - | - | - | 35 | 0.91 | 0.614 |
| | Residuals | 137 | - | - | 102 | - | - |
| Number of seeds in 10 fruits | Group | 4 | 152.54 | **<2.2e-16** | 4 | 170.78 | **<2.2e-16** |
| | Tray | - | - | - | 35 | 1.59 | **0.0367** |
| | Residuals | 146 | - | - | 111 | - | - |

| | | **Generalized linear models (quasibinomial, link="logit"), ANOVA (type III tests)** | | | | | |
|---|---|---|---|---|---|---|---|
| | | **~ Group** | | | **~ Group + Tray_ID** | | |
| | | **df** | **F** | **p-value** | **df** | **F** | **p-value** |

| | | | | | | | |
|---|---|---|---|---|---|---|---|
| Pollen viability | Group | 4 | 24.39 | **2.97e-15** | 4 | 21.06 | **1.12e-12** |
| | Tray | - | - | - | 35 | 0.723 | 0.862 |
| | Residuals | 137 | - | - | 102 | - | - |
| Proportion of normal seeds | Group | 4 | 59.16 | **<2.2e-16** | 4 | 83.72 | **<2.2e-16** |
| | Tray | - | - | - | 35 | 1.35 | 0.121 |
| | Residuals | 146 | - | - | 111 | - | - |

-: Not applicable. p-values < 0.05 were highlighted in bold.

**Figure 4–Source Data 1** Additive and non-additive gene expression in allotetraploid groups

| | | Additive Expression* | | Nonadditive expression | | | | | | | |
|---|---|---|---|---|---|---|---|---|---|---|---|
| | | | | Expression-level dominance (ELD) | | | | Transgressive expression (TRE) | | | |
| | | | | Cg-ELD | | Co-ELD | | Over-TRE | | Under-TRE | |
| | Expression category | a | b | c | d | e | f | g | h | i | j |
| | Plant group | 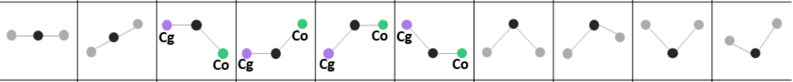 | | | | | | | | | |
| Flower | Sd | 18686 (86.3%) | 1542 (7.1%) | 821 (3.8%) | 85 (0.4%) | 496 (2.3%) | 16 (0.1%) | 1 (0.0%) | 0 | 0 | 0 |
| | Sh | 18711 (86.4%) | 1583 (7.3%) | 812 (3.8%) | 56 (0.3%) | 448 (2.1%) | 32 (0.2%) | 4 (0.0%) | 0 | 1 (0.0%) | 0 |
| | Cbp | 18021 (83.2%) | 1302 (6.0%) | 797 (3.7%) | 141 (0.7%) | 769 (3.6%) | 332 (1.5%) | 222 (1.0%) | 30 (0.1%) | 27 (0.1%) | 6 (0.0%) |
| Leaf | Sd | 16522 (88.1%) | 1374 (7.3%) | 549 (2.9%) | 9 (0.1%) | 256 (1.4%) | 45 (0.2%) | 3 (0.0%) | 0 | 0 | 0 |
| | Sh | 16520 (88.1%) | 1384 (7.4%) | 513 (2.7%) | 29 (0.2%) | 234 (1.3%) | 78 (0.4%) | 0 | 0 | 0 | 0 |
| | Cbp | 15894 (84.7%) | 925 (4.9%) | 667 (3.6%) | 187 (1.0%) | 476 (2.5%) | 363 (1.9%) | 189 (1.0%) | 25 (0.1%) | 22 (0.1%) | 10 (0.1%) |

*Partial expression-level dominance (expression level in allotetraploids was not the same as the mid-parent value but still in the middle of the two diploid groups) was included in category b of additive expression. The ten gene expression categories were: a) additive expression with no parental differentiation, b) partial ELD or additive expression with parental differentiation, c) Up-regulated ELD toward diploid *C. grandiflora* (Cg2), d) Down-regulated ELD toward Cg2, e) Up-regulated ELD toward diploid *C. orientalis* (Cg2), f) Down-regulated ELD toward Co2, g) Up-regulated TRE with no parental differentiation, h) Up-regulated TRE with parental differentiation, i) Down-regulated TRE with no parental differentiation, g) Down-regulated TRE with parental differentiation.

**Figure 8–Source Data 1** Expression level fold-change (log2FC) of homoeologs relative to the corresponding gene in diploid groups among genes with expression level dominance (ELD) in flowers or leaves.

| Tissue | Source of ELD | log2FC of EL-dominant homoeologs (mean±se) | log2FC of EL-recessive homoeologs (mean±se) | p-value* |
|---|---|---|---|---|
| Flower | Sd/Sh ELD | 0.639±0.013 | 1.823±0.034 | <2.2e-16 |
| | Cbp-specific ELD | 1.208±0.037 | 2.621±0.058 | <2.2e-16 |
| Leaf | Sd/Sh ELD | 0.792±0.024 | 1.933±0.048 | <2.2e-16 |
| | Cbp-specific ELD | 1.439±0.045 | 3.067±0.072 | <2.2e-16 |

*The difference of expression fold change between EL-dominant and EL-recessive homoeologs was tested by Welch's two sample t-tests.

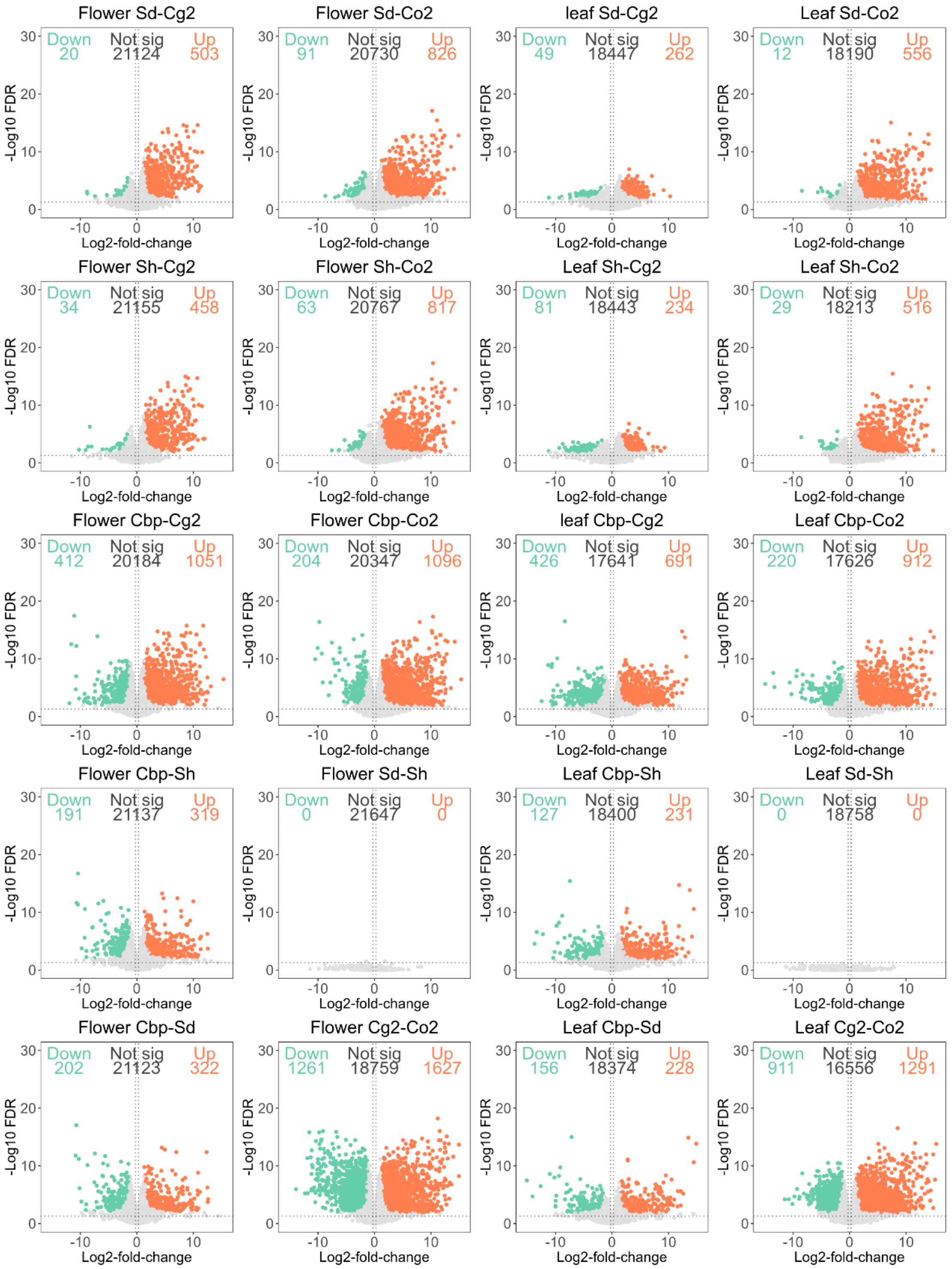


**Supplementary file 3** Differentially expressed genes (DEGs) in pair-wise contrasts among the five *Capsella* plant groups in flowers and leaves. Genes with expression (CPM > 1) in at least two samples were used for the differential expression (DE) analysis. DE analysis was conducted with the R package edgeR (version 3.28.1; Robinson *et al.*, 2010), using TMM normalized unphased gene expression levels. Genes with fold-change > 2 and false discovery rate < 0.05 were considered significant DEGs.

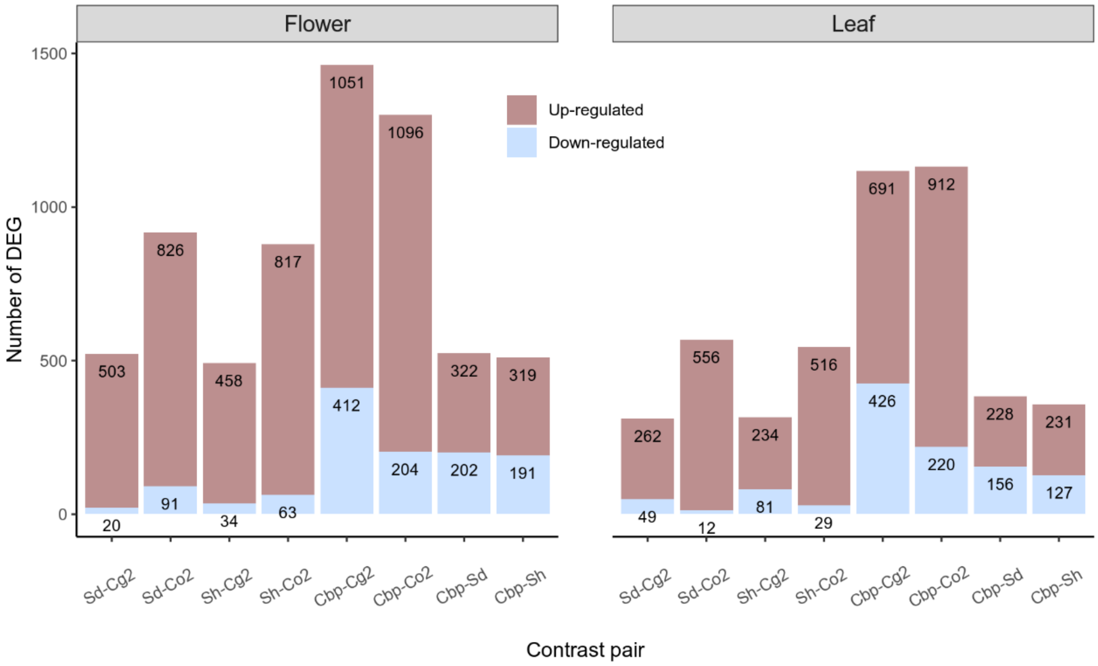


**Supplementary file 4** Summary of differential expression analyses of allotetraploid groups. The number of up-regulated and down-regulated differentially expressed genes (DEGs) in each group pair were annotated on plots. DE analysis was performed on 21,937 genes in flowers and 18,999 genes in leaves, with a threshold of $FC > 2$ and $FDR < 0.05$. Gene expression levels were normalized with the trimmed mean of M-values (TMM) method.
